# Transcranial Functional Ultrasound Imaging Detects Focused Ultrasound Neuromodulation Induced Hemodynamic Changes *In Vivo*

**DOI:** 10.1101/2024.03.08.583971

**Authors:** Jonas Bendig, Christian Aurup, Samuel G. Blackman, Erica P. McCune, Seongyeon Kim, Elisa E. Konofagou

## Abstract

**Background:** Focused ultrasound (FUS) is an emerging non-invasive technique for neuromodulation in the central nervous system (CNS). Functional ultrasound imaging (fUSI) leverages ultrafast Power Doppler Imaging (PDI) to detect changes in cerebral blood volume (CBV), which correlate well with neuronal activity and thus hold promise to monitor brain responses to FUS.

**Objective:** Investigate the immediate and short-term effects of transcranial FUS neuromodulation in the brain with fUSI by characterizing hemodynamic responses.

**Methods:** We designed a setup that aligns a FUS transducer with a linear array to allow immediate subsequent monitoring of the hemodynamic response with fUSI during and after FUS neuromodulation (FUS-fUSI) in lightly anesthetized mice. We investigated the effects of varying pressures and transducer positions on the hemodynamic responses.

**Results:** We found that higher FUS pressures increase the size of the activated brain area, as well as the magnitude of change in CBV and could show that sham sonications did not produce hemodynamic responses. Unilateral sonications resulted in bilateral hemodynamic changes with a significantly stronger response on the ipsilateral side. FUS neuromodulation in mice with a cranial window showed distinct activation patterns that were frequency-dependent and different from the activation patterns observed in the transcranial model.

**Conclusion:** fUSI is hereby shown capable of transcranially monitoring online and short-term hemodynamic effects in the brain during and following FUS neuromodulation.

## Introduction

Focused ultrasound (FUS) has shown the ability to noninvasively modulate neuronal activity in the central nervous system of different animal species as well humans [1–9]. In contrast to established neuromodulatory techniques like deep brain stimulation, transcranial magnetic stimulation or transcranial (direct) current simulation, FUS combines a favorable safety profile with the ability to target deep brain structures with spatial resolution in the millimeter (e.g. humans) or sub-millimeter range (e.g. rodents) [9,10]. In a previous study, we introduced a robust technique for performing FUS neuromodulation in mice *in vivo* [11]. However, *in silico* ultrasound simulations indicated that transcranial pressure field patterns are difficult to predict and that intracranial acoustic reverberations could generate additional pressure peaks sufficient to activate the brain outside of the intended focal target. More direct measurements of evoked brain activity are therefore needed to fully assess the acute and long-term effects of FUS in the targeted area as well as connected brain regions.

Electroencephalography (EEG) [12,13] and functional magnetic resonance imaging (fMRI) [14,15] are the most common techniques for studying neuronal activity in animal models and humans. However, EEG is not capable of directly localizing activity in deep brain regions and fMRI requires long imaging sessions in a spatially confined MRI scanner with high capital costs [16–18]. In contrast, functional ultrasound imaging (fUSI) is an emerging imaging technique [19–21] that allows monitoring of stimulus-evoked activity and functional connectivity in the whole brain with a comparatively small ultrasound array [22–25]. fUSI uses ultrafast Power Doppler Imaging (PDI) to measure changes in cerebral blood volume while suppressing signals from the surrounding tissue through the implementation of advanced spatiotemporal filtering techniques such as singular value decomposition (SVD) [26,27]. Analogous to fMRI, fUSI leverages neurovascular coupling and has been shown to correlate well with neuronal activity and local field potentials [28,29]. The spatial resolution is similar to that of fMRI [30] but it attains greater temporal resolution [16].

The principal challenge in applying transcranial fUSI to the brain is the substantial acoustic attenuation induced by the skull. Consequently, most implementations of fUSI to date have relied on the removal or thinning of the skull bone [18]. Previous work by our group and others has demonstrated transcranial applications of fUSI for detecting hemodynamic changes in the mouse brain [23,31–33], allowing for a fully noninvasive ultrasound-based functional brain imaging technique. Achieving transcranial fUSI in combination with FUS could allow noninvasive neuromodulation with simultaneous monitoring of neuromodulatory effects in a large field of view.

In this study, we developed a simultaneous FUS and power Doppler imaging transducer configuration to assess the immediate and short-term effects of FUS neuromodulation with transcranial conventional fUSI. We demonstrate that the size of the activated area and the change in cerebral blood volume (CBV) are positively correlated with the magnitude of the applied pressure and that unilateral sonications result in bilateral hemodynamic changes with a significantly stronger response on the ipsilateral side. FUS neuromodulation in mice with a cranial window shows distinct activation patterns that are frequency-dependent and different from the activation patterns observed in the transcranial model.

## Methods

### Animal preparation

Young male wild-type mice between 8 to 12 weeks of age (C57BL/6, n = 22) were used for transcranial experiments in this study. To assess possible effects of the skull, a subgroup of mice between 8 to 20 weeks of age (C57BL/6, n = 5) was implanted with a chronic cranial window covered by a polymethyl pentene membrane as described by Brunner et al [20]. Mice implanted with a cranial window were allowed to rest for 2 weeks before experiments were performed. Anesthesia was induced with isoflurane (1-3 %) and supplementary oxygen (0.8 L/min). The absence of a pedal reflex confirmed induction and isoflurane was then decreased and adjusted between 0.5 – 1 % to maintain light anesthesia without producing gasping from low oxygenation, to reduce motion artifacts. The subject’s head was fixed by a stereotactic frame (Model 900, David Kopf Instruments, Tujunga, CA, USA) and elastic bands were passed over the body to mitigate motion artifacts. The animal’s head was shaved and depilatory cream was used to remove all fur before acoustic gel was placed on the animal’s head. Mice with cranial windows were fixed with a 3D-printed holder that connected to the implanted headpost and acoustic gel was placed on the membrane covering the cranial window. A data acquisition system (MP150, Biopac Systems Inc, Goleta, CA) was used to acquire pulse-oximetry signals (MouseOx+, Starr Life Sciences, Oakmont, PA, USA) recorded from a sensor placed on the shaved thigh. The pulse-oximeter was used in conjunction with intermittent toe pinches to monitor the depth of anesthesia. The ideal depth of anesthesia during experiments corresponded with heart rates above 400 bpm and an unresponsive pedal reflex.

### FUS Neuromodulation

We used single-element spherical segment annular focused ultrasound transducers (H-204 and H-215, Sonic Concepts Inc, Bothell, WA, USA) that were confocally aligned with an ultrasound imaging array (L22-14vXLF, Vermon S.A., Tours, France). Each transducer had an acoustic coupling cone attached to the transducer face with an acoustically transparent membrane (Tegaderm, 3M Company, Maplewood, MN, USA) placed over its opening. The sealed coupling chamber was filled with deionized water and degassed using a degassing system (WDS105+; Sonic Concepts, Bothell, WA, USA). FUS transducers were calibrated in a degassed water tank using a hydrophone (HGL0200, Onda Corporation, Sunnyvale, CA, USA) in free field or with ex vivo skulls for estimating attenuation and reporting derated pressures. FUS sequences were driven by a function generator and amplified with an RF amplifier (325LA, Electronics Innovation Ltd., Rochester, NY, USA). A schematic of the setup is depicted in Figure 1 a and the acoustic parameters and focal size of the transducers are listed in Table 1. For targeting, the confocal FUS-fUSI transducer system was lowered into the acoustic gel on the scalp, ensuring that no bubbles were trapped along the beam path. B-mode imaging was used to verify adequate acoustic coupling. Targeting was performed by first landmarking the interaural line using B-mode imaging to locate highly reflective metal syringe tips temporarily placed on the stereotactic ear bars (i.e. interaural line). The syringe tips were then removed, taking care not to leave trapped air bubbles that could result in signal loss or artifacts. Neuronavigation could then be performed by manually translating the transducer system in the anteroposterior or mediolateral (ML) directions according to a reference atlas [34] using the micromanipulator (David Kopf Instruments, Tujunga, CA, USA). In animals with cranial windows, targeting was performed by using the anterior border of the cranial window (Bregma +2 in the anteroposterior direction) as a reference point and moving the transducer system with a motorized stage. We used post-hoc mapping of the power Doppler images based on custom vasculature reference scans relative to Bregma over the whole brains of two mice (0.1 mm steps) and found that the imaging planes used in this work span between -0.2 mm to -2.0 mm from Bregma in the anteroposterior direction. Imaging planes of all figures shown and the median (range) of imaging planes used in datasets can be found in the figure captions. Sonications were performed along the midline (± 0 mm ML) or at ± 1 mm ML to investigate lateralization effects. To mitigate auditory artifacts associated with pulsed ultrasound, we implemented an amplitude-modulated (AM) sequence [35,36]. The amplitude modulation was based on a sine wave (half cycle) and the energy deposited in the tissue was equal to a duty cycle of 64 %. A block design was implemented consisting of a 60 s baseline imaging block followed by four 20 s FUS blocks with intervals randomized between 60 and 90 s. During the FUS blocks, the function generator output was triggered by a programmable research ultrasound system (Vantage 256, Verasonics, Kirkland, WA) directly after image acquisition to avoid any interference between FUS and fUSI (Figure 1 b). Supplementary Video 1 shows a sequence where FUS and fUSI were not interleaved, which led to pronounced FUS-induced artifacts. Comparable artifacts were not observed with the interleaved sequence employed in this study.

**Figure 1.**
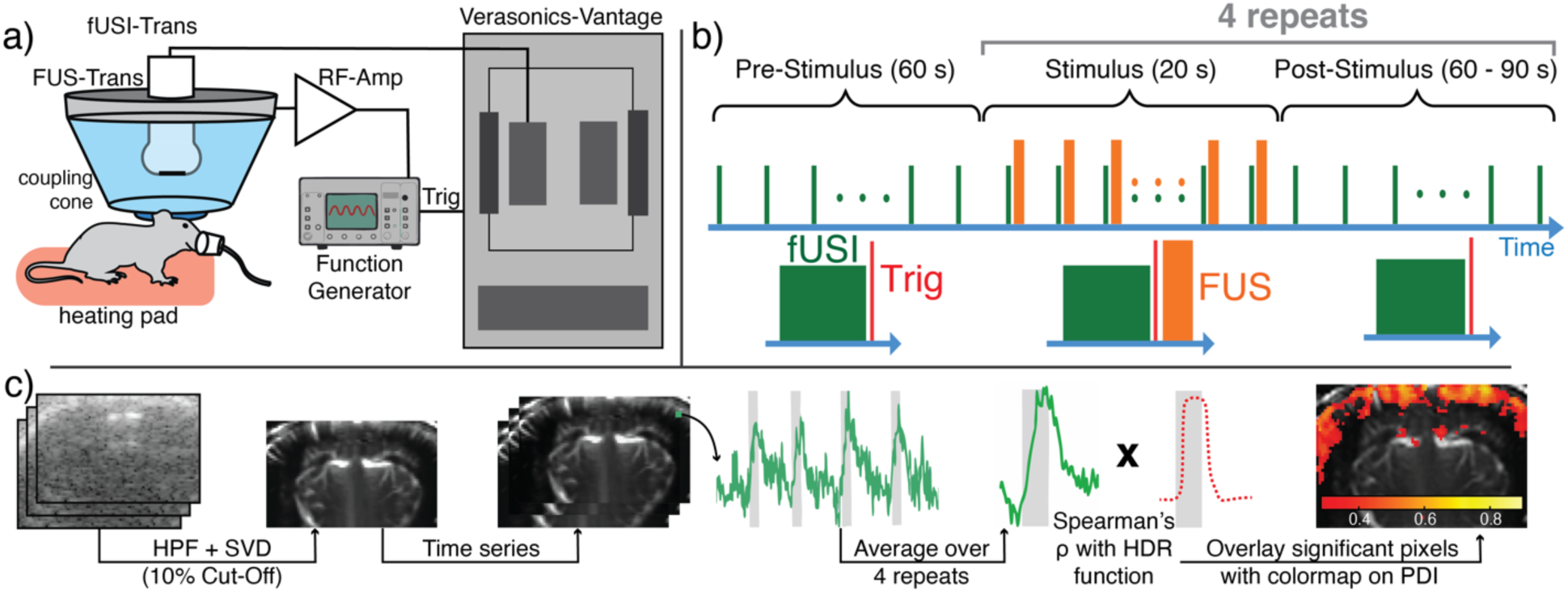
Experimental setup **(a)**, fUSI and FUS stimulus block design **(b)** and fUSI data processing pipeline **(c)**. Abbreviations: FUS – focused ultrasound, fUSI – functional ultrasound, HDR – hemodynamic response, HPF – high pass filter, PDI – power doppler image, RF-Amp – radiofrequency amplifier, SVD – singular value decomposition, Trig – Trigger.

**Table 1.** FUS and fUSI equipment and acoustic parameters.

| **FUS** |  |
| --- | --- |
| **Focal Size (-6 dB)** | H-204: 0.8 mm (lateral), 8 mm (axial)<br>H-215: 0.3 mm (lateral), 2 mm (axial) |
| **Acoustic Parameters** | Carrier Frequency:<br>H-204: 1.68 MHz (3<sup>rd</sup> Harmonic)<br>H-215: 4 MHz<br>Amplitude Modulation Freq: 1 kHz<br>Burst Duration: 150 ms<br>Sonication Frequency:<br>H-204: 1 Hz<br>H-215: 2 Hz<br>Peak Negative Pressures:<br>0.8-2.6 MPa (Derated, transcranial, H204)<br>0.7-2.8 MPa (Cranial window, H204)<br>1.4-3.6 MPa (Cranial window, H215) |
| **fUSI** |  |
| **Acoustic Parameters** | Center frequency: 15.0 MHz<br>Compounded Plane Wave Imaging<br>Plane Angles: 5 (-6° to 6°)<br>Compounded Image Rate: 500 Hz<br>Compounded Images in PDI:<br>200 (transcranially)<br>70 (craniotomized) |
| **Filtering** | Singular Value Decomposition: 10% of singular values removed<br>High-Pass Filter: Butterworth (2<sup>nd</sup> order), 8-Hz cutoff frequency |

### Sham conditions and safety monitoring

To rule out activations unrelated to FUS or connected to auditory artifacts, two sham conditions were implemented. In the first sham condition, the amplifier was turned off and in the second sham condition a 110 dB, 10 kHz tone (square wave) was delivered instead of the FUS stimulus in the same temporal pattern. The tone was generated with an Arduino microcontroller coupled to a speaker (BA735 Digital, Boston Acoustics, Woburn, MA) that was placed 10 cm in front of the mouse’s head. Speaker volume was measured at 10 cm distance at the position of the mouse head with the National Institute for Occupational Safety and Health Sound Level Meter App.

To investigate the safety of the FUS parameters, 2 mice were transcranially sonicated with the highest pressures used in this study (1.68 MHz, 2.6 MPa) and the remaining parameters as described in Table 1. Mice were sacrificed 24 hours after the end of the sonication, and brain tissue in the focal plane was stained with H&E for histopathological investigation of tissue morphology and possible blood extravasation. In addition, transcranial temperature measurements were performed in 2 separate mice in two locations by placing a thermocouple (HYP1, T-type needle thermocouple, 0.25 mm diameter, Omega Engineering, Norwalk, CT) directly under the skull or at the estimated location of the center of the FUS focus in the brain with a micromanipulator. At each location, the thermal response was maximized by micromanipulation of the focus in the anteroposterior and mediolateral direction in 0.1 mm steps. FUS pulses with 1.68 MHz were delivered with 0.8, 1.7 and 2.6 MPa for a total duration of 20 s with the parameters described in Table 1. Temperature measurements were corrected for the viscous heating artifact by iterative curve fitting of the Bioheat Transfer Equation as described previously [37].

### Functional Ultrasound Imaging (fUSI)

fUSI was implemented to detect and characterize stimulus-evoked hemodynamic changes in the mouse brain. The imaging sequence was generated using a programmable research ultrasound system to perform ultrafast compounded plane wave imaging. fUSI was performed by acquiring a time series of coronal power Doppler images (PDI) at a resolution of 103 μm, assuming a sound speed of 1546 m/s for brain tissue. Array specifications and imaging parameters utilized are provided in Table 1. A single compounded image was generated by averaging delay-and-sum reconstructed ultrasound images acquired from multiple plane wave transmits. A PDI was generated by first applying a high-pass filter to a stack of compounded images followed by spatiotemporal filtering using singular value decomposition (SVD), where the upper 10 % of eigenvalues were discarded to filter out tissue clutter signal [26,27]. SVD filtering cannot remove the influence of large motion artifacts, so outlier frames were removed whose mean image value was three standard deviations above the mean image value of the remaining image set. The pixel intensity data, representing CBV, was averaged across the four stimuli before performing the statistical analysis. The image processing steps are outlined in Figure 1 c.

### Correlation Analysis

Statistical analyses were performed to identify pixels exhibiting intensity time courses that were significantly correlated with applied stimulus patterns as described previously [31]. The binary stimulus vector (i.e. FUS ON vs FUS OFF) was convolved with a modified hemodynamic response function (HRF) [20] to generate a more physiologically relevant HRF regressor for computing correlation coefficients. The following parameters were used: delay of response (relative to onset) = 4 s, delay of undershoot (relative to onset) = 36 s, dispersion of response = 0.5, dispersion of undershoot = 1, ratio of response to undershoot = 20, onset = 0 s, length of kernel (seconds) = 16 s. Pixel-wise Spearman correlation coefficients were then computed between the regressor and PDI time series. Pixels were defined as significantly activated with ρ > 0.29 (transcranial experiments) or ρ > 0.19 (craniotomized experiments). The thresholds were obtained as described by Brunner et al. [20] by using Fisher’s transform with Z = 2.58 (p < 0.01) for 80 or 175 time points, respectively. Significantly activated pixels were used to create binary maps. The computed correlation values were then remapped using the binary maps to be overlaid onto the mean PDI (Figure 1c). Correlation maps were filtered in space with a median filter (kernel size = 3) and CBV response curves were filtered with a rolling mean (window size = 3) before being displayed.

## Results

### fUSI transcranially detects hemodynamic responses during and following FUS

To investigate if FUS can induce a hemodynamic response in the mouse brain, we compared active FUS with two sham conditions: Sham trials performed with the amplifier powered off showed no significant activation (n = 22 trials in 6 mice, transcranially, Figure 2 c/d). Similarly, an auditory sham condition with a 110 dB loud 10 kHz tone (n = 3 mice implanted with a cranial window, Figure 2 e/f) showed no activation. However, FUS routinely produced widespread hemodynamic responses in sonicated subjects in the transcranial (p < 0.0001 vs sham, Wilcoxon signed-rank test) and cranial window conditions. Activity was typically observed both within the focal region and across the cortex in both hemispheres.

**Figure 2.**
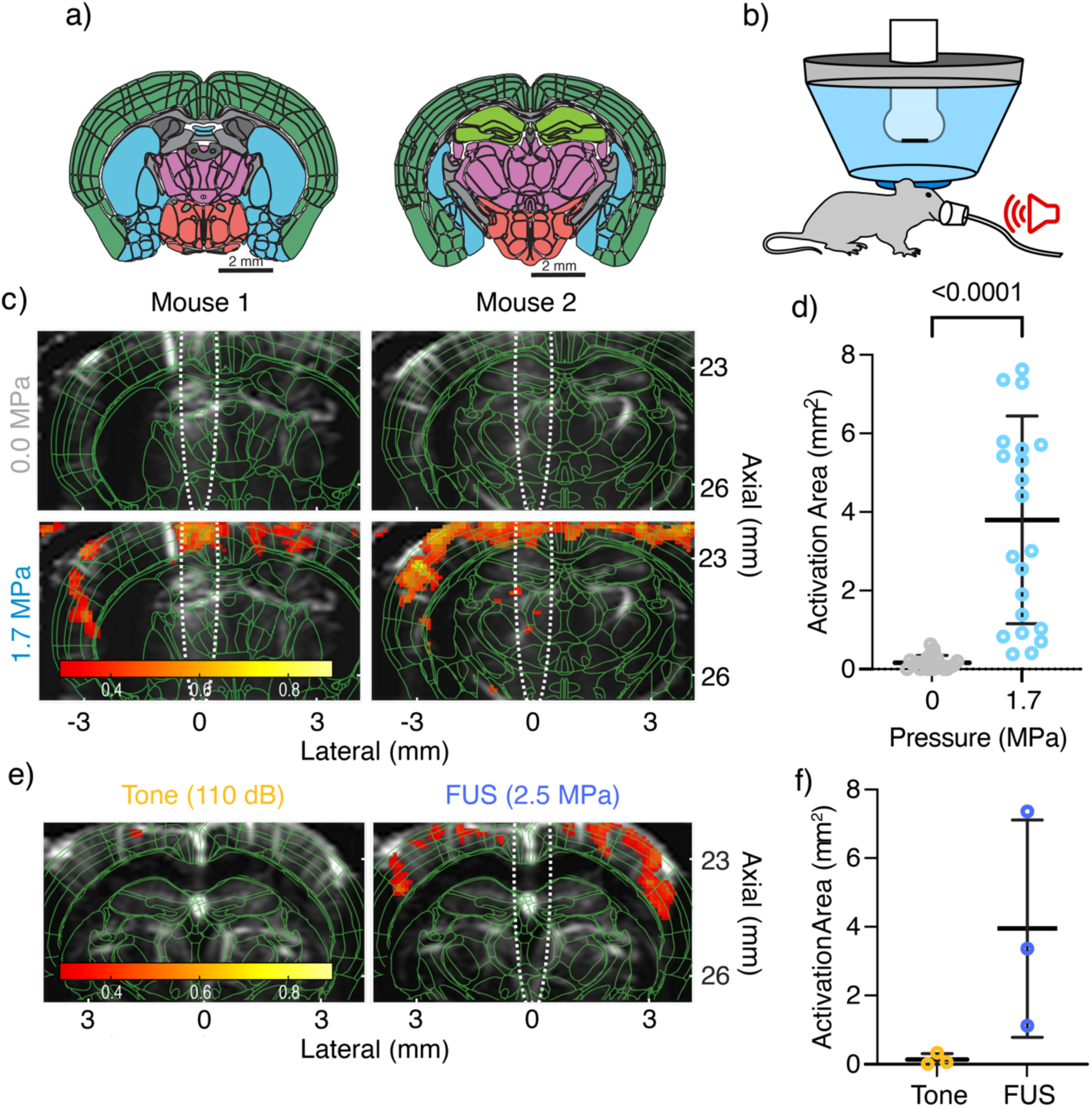
Hemodynamic response in FUS and sham trials. **(a)** Schematic representation of the imaging planes in **b** (left: Mouse 1, AP Bregma – 0.6 mm; right: Mouse 2, AP Bregma – 1.4 mm) and c (right, AP Bregma – 1.4 mm) and. Brain regions are colored analogous to the Allen Mouse Brain Atlas [38] (Cortex: Dark green, Cerebral nuclei: Blue, Hippocampus: Light green, Thalamic nuclei: Purple, Hypothalamic nuclei: Red, Fiber tracts: Light gray, Ventricular System: Dark gray). **(b)** Schematic representation of the setup used for sham/active trials (Functional ultrasound transducer: White, FUS transducer: Gray, Water-filled coupling cone: Light blue, Coupling gel: Dark blue, Speaker: Red). **(c)** Activity maps of two subjects receiving FUS-Off sham and active FUS conditions overlaid onto PDI (grayscale) and brain regions (green). The predicted acoustic focus assessed during transducer calibration is overlaid with a white dotted line. **(d)** Wilcoxon matched-pairs signed rank test revealed a significantly greater activation area in the FUS group (n = 22 paired trials in 6 mice, median AP direction of imaging planes: -0.75 mm from Bregma, Range = - 0.5 mm to -1.4 mm). **(e)** Representative activity maps for a subject receiving auditory stimulation only (tone) and active FUS overlaid onto PDI (grayscale) and brain regions (green). **(f)** Comparison of activation areas for auditory stimulation and active FUS condition (n = 3 paired trials in 3 mice, median AP direction of imaging planes: -1.5 mm from Bregma, Range = -1.4 mm to -1.6 mm). Data in panels c/d is based on transcranial imaging, while data in e/f was obtained in mice with cranial windows. Graphs depict individual observations (circles) with mean (central line) and standard deviations (whiskers). Activity maps and color bars show Spearman correlation coefficients. Only significantly correlated pixels (Spearman’s ρ > 0.285 are displayed)

### Hemodynamic responses in the brain depend on FUS pressure

To determine whether response patterns depend on the applied acoustic pressure, 4 subjects were sonicated with 3 different FUS pressures in randomized order for a total of 10 trials (n = 3 trials for 2 subjects, n = 2 trials for 1 subject and n = 1 trial for 2 subjects) and hemodynamic changes were detected with transcranial fUSI. A sample of pressure conditions for 3 subjects is provided in Figure 3 a.

**Figure 3.**
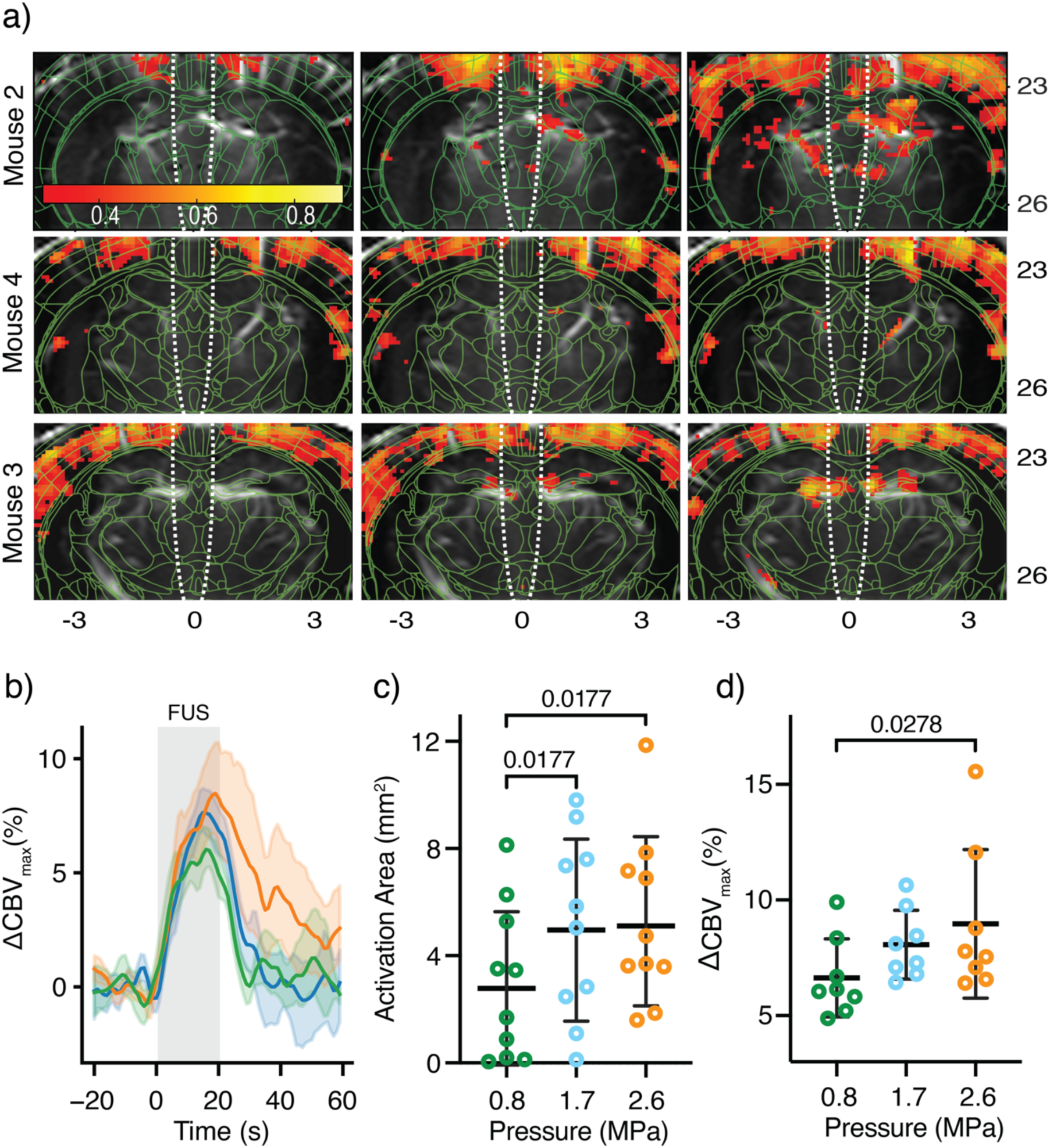
Higher pressures of transcranial FUS induce greater activation in mice. **(a)** Activity maps of significantly correlated pixels (Spearman’s ρ > 0.285) overlaid onto PDI (grayscale) and brain regions (green) for 3 subjects receiving 0.8, 1.7, and 2.6 MPa FUS. Predicted acoustic foci (white, dotted line) assessed during transducer calibration are overlaid onto the images. Imaging planes: Upper row AP Bregma -0.5 mm, middle row AP Bregma -1.0 mm, bottom row AP Bregma -1.5 mm **(b)** Mean CBV changes (solid) and 95 % confidence interval for all jointly significantly correlated pixels across all trials (n = 8 paired trials in 4 mice, no pixels found in 2 trials). Multiple Wilcoxon matched-pairs signed rank tests (with Bonferroni correction, after significant Friedman tests: p = 0.0063 **(c)** and p = 0.0303 **(d)**) revealed significantly greater activation area **(c)** and peak CBV changes in jointly significant pixels **(d)** as pressure increased (n = 10 paired trials in 4 mice, no jointly significant pixels in 2 trials, median AP direction of imaging planes: -1.0 mm from Bregma, Range = -0.2 mm to -1.5 mm). Data is depicted as individual observations (circles) with mean (central line) and standard deviations (whiskers). Activity maps and color bars show Spearman correlation coefficients.

We found a significant difference between the activation area for different pressure groups (p = 0.0063, Friedman test) with higher pressures of 1.7 and 2.6 MPa activating a significantly greater area compared to the 0.8 MPa group (p = 0.0177 for both, Wilcoxon matched-pairs signed rank tests with Bonferroni correction, Figure 3 c). To further investigate the effects of pressure on the CBV response, we calculated the mean CBV change over time in all jointly significantly correlated pixels that were common to each of the three pressure conditions in individual trials (2 trials were omitted from the analysis because the 0.8 MPa condition did not yield any significance). We found a significantly higher maximal CBV increase in the 2.6 MPa group compared to the 0.8 MPa group (p = 0.0278, Wilcoxon matched-pairs signed rank test with Bonferroni correction, after significant Friedman test (p = 0.0303), Figure 3 d).

To investigate possible effects of the skull on the response patterns, FUS was delivered with different pressures in 5 mice implanted with cranial windows. We further evaluated possible effects of frequency and focal size by using a 4 MHz transducer to deliver FUS with comparable pressures. Like the transcranial condition, there were significant differences between the pressure groups for both tested frequencies (Friedman test, 1.68 MHz: p = 0.0010, 4.0 MHz: p < 0.0001). A Dunn’s test corrected for multiple comparisons was performed as a post-hoc analysis to account for the small group size. We found significantly larger activated areas in 2.8 MPa FUS compared to the lower pressures in the 1.68 MHz condition (Figure 4 b). Similarly, with 4 MHz FUS the activated area with pressures ≥ 3.0 MPa was significantly larger compared to 1.4 MPa or 1.8 MPa (Figure 4 c). Clear trends for an increase in activated areas start at pressures of 2.1 MPa in the 1.68 MHz condition and at 2.3 MPa in the 4 MHz condition (Figure 4 b/c). Compared to transcranial FUS there was no activation in lower pressure conditions (i.e. 0.8 MPa and 1.7 MPa). Especially in lower pressures, cortical activation was less pronounced in mice with cranial windows, while subcortical responses appeared enhanced (Figure 4 a). Interestingly, we found significant negative correlations in the cranial window condition (Figure 4 a, blue colored regions) while they were absent in the transcranial condition (no difference for negatively correlated pixels compared to sham sonication, no negative correlations in different pressure conditions). Negative correlations appeared more pronounced with higher frequency but showed a significant effect of pressure on the area of significant negatively correlated pixels in both frequency conditions (1.68 MHz: Friedman test = 0.0104; 4.0 MHz: Friedman test = 0.0003, Supplementary Figure 3).

**Figure 4.**
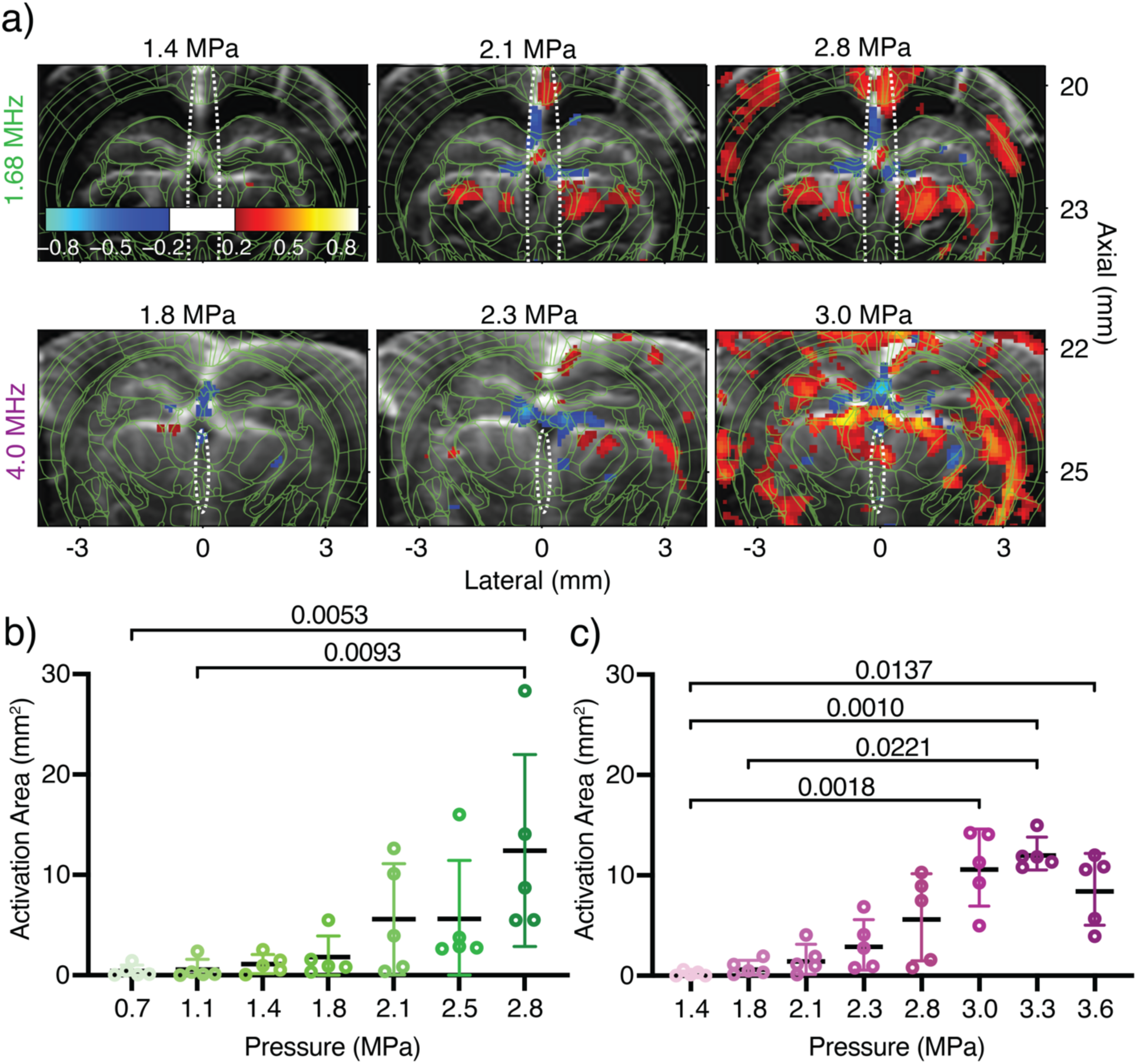
Higher pressures with different center frequencies induce greater activation in craniotomized mice. **(a)** Representative activity maps of significantly positive correlated pixels (Spearman’s ρ > 0.19, red hues) and significantly negative correlated pixels (Spearman’s ρ < -0.19, blue hues) for 2 subjects receiving FUS with 1.68 MHz or 4.0 MHz center frequency under different pressure conditions. The predicted acoustic foci (white) assessed during transducer calibration and brain regions (green) are overlaid onto the PDIs. Imaging planes: Upper row AP Bregma -1.4 mm, bottom row AP Bregma -1.7 mm. **(b)** Activation area (positively correlated pixels) for different pressures in the 1.68 MHz condition (n = 5 paired trials in 5 mice, median AP direction of imaging planes: -1.5 mm from Bregma, Range = -1.4 mm to -1.9 mm; Friedman test: p = 0.0010). **(c)** Activation area (positively correlated pixels) for different pressures in the 4.0 MHz condition (n = 5 paired trials in 5 mice, median AP direction of imaging planes: -1.5 mm from Bregma, Range = -1.4 mm to -2.0 mm; Friedman test: p < 0.0001). Areas of negatively correlated pixels are depicted in Supplementary Figure 2. Data is depicted as individual observations (circles) with mean (central line) and standard deviations (whiskers). Activity maps and color bars show Spearman correlation coefficients. Dunn’s test (corrected for multiple comparisons) was used for post-hoc comparisons after significant Friedman tests.

### Transcranial FUS induces hemodynamic responses with ipsilateral bias

To investigate if FUS causes lateralized responses, sonications were targeted 1 mm left and right of the midline in each imaging plane in 11 paired trials across 6 subjects. Unilateral sonications produced mostly cortical bilateral hemodynamic responses (Figure 5 a) as shown before, however, significantly greater activation areas were observed in the ipsilateral sonicated hemisphere (Figure 9 b). In addition, the mean Spearman’s ρ was higher in the hemispheres where FUS was delivered compared to the contralateral hemispheres (Figure 5 c).

**Figure 5.**
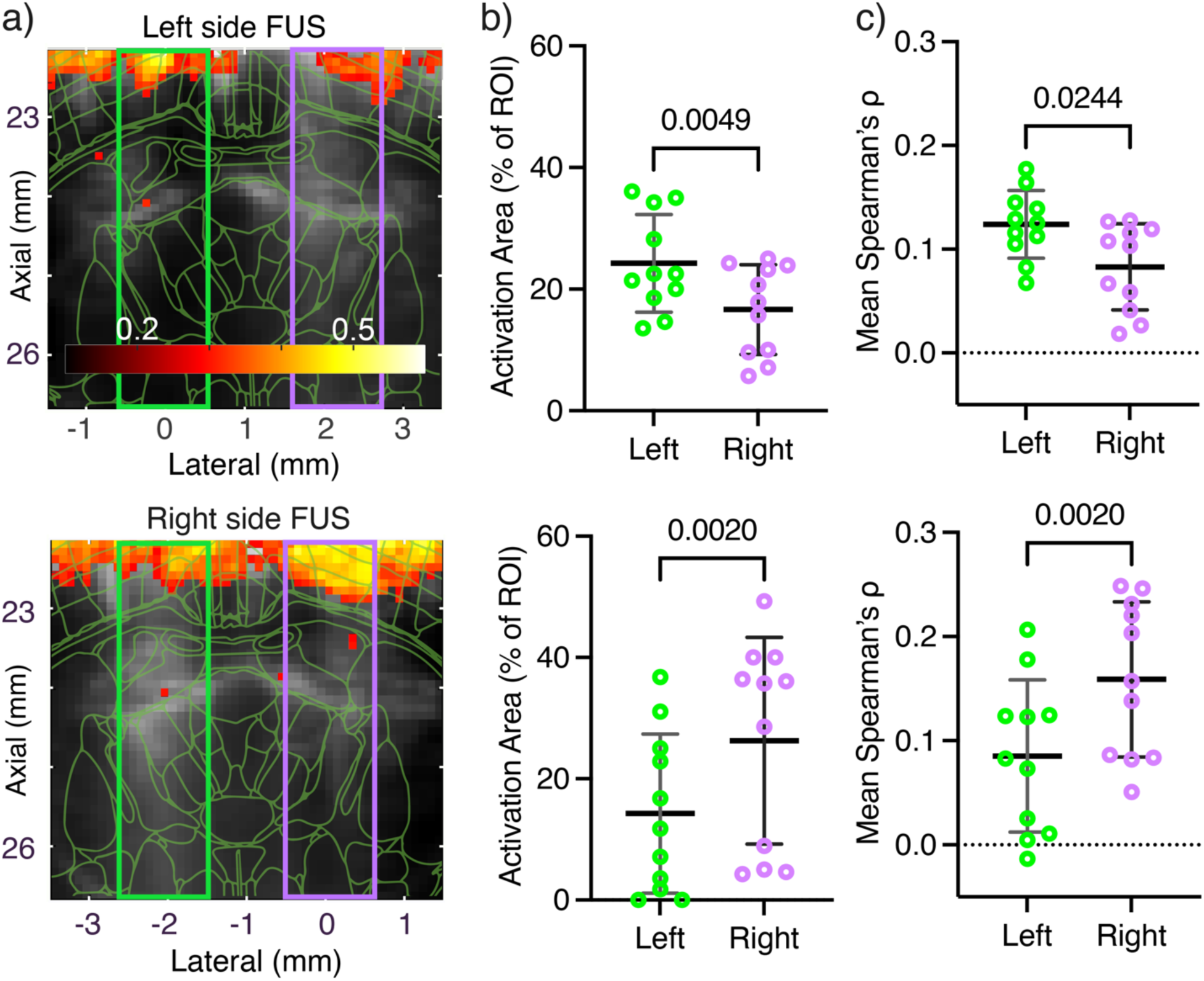
Transcranial FUS shows ipsilateral bias in CBV responses to unilateral sonications. Averaged activation maps of significantly correlated pixels (Spearman’s ρ > 0.285) overlaid onto averaged PDI (grayscale) and brain regions of median imaging plane (green) for left **(a**, upper row) and right **(a**, lower row) hemispheric sonications (n =11 paired trials in 6 mice, median AP direction of imaging planes: -0.5 mm from Bregma, Range = -0.4 mm to -1.3 mm). Rectangles show regions of interest (ROI) in the right (purple) and left (green) hemispheres that were used for analyses in **b** and **c**. The dotted white line shows the predicted acoustic foci (white) assessed during transducer calibration. FUS induces significantly more activated pixels **(b)** and higher mean correlation coefficients **(c)** in the sonicated ipsilateral ROI compared to the unsonicated contralateral ROI. Data is depicted as individual observations (circles) with mean (central line) and standard deviations (whiskers). Activity maps and color bars show Spearman correlation coefficients. Wilcoxon matched-pairs signed rank test was used for group comparisons.

### Safety of transcranial FUS-fUSI

We found no signs of structural changes or blood cell extravasations in H&E staining in n = 2 mice that were sacrificed 24 h following FUS with the highest pressure used for transcranial FUS (center frequency = 1.68 MHz, PNP = 2.6 MPa, Supplementary Figure 3 c/d). Thermocouple measurements (n = 2 mice per target, center frequency = 1.68 MHz) showed maximal temperature increases of 0.81 °C, 1.44 °C or 3.44 °C at –2 mm DV in the brain tissue and 0.70 °C, 2.08 °C or 3.58 °C directly under the skull for derated peak negative pressures of 0.8 MPa, 1.7 MPa and 2.6 MPa, respectively (Supplementary Figure 3 a/b).

## Discussion

This study presents the first implementation of transcranial fUSI in combination with FUS to investigate neuromodulatory effects in the CNS by analyzing changes in CBV. We demonstrate that higher pressures significantly increase the activated area in the brain and induce stronger CBV increases, while lateralized sonications result in CBV responses with ipsilateral bias in mice.

Our results demonstrate robust and repeatable bilateral activation of cortical and subcortical areas during FUS neuromodulation not strictly limited to the focal area. This finding is seemingly in contrast with several studies demonstrating focal activation and specific behavioral responses during or following FUS [2,5,39–42]. However, our group and others have demonstrated off-target activations likely associated with brain regions connected to the same network as the targeted areas [43,44] and network-associated changes in functional connectivity following FUS neuromodulation are widely reported [45–48]. The long neuromodulation periods (20 s) and high pressures (up to 3.6 MPa) employed in this study make it conceivable that connected brain areas were activated, while the comparatively low frame rate of fUSI (1 – 2 Hz) did not allow the identification of a focal starting point. This notion is partially supported by the facts that the induced CBV responses remained consistent at single transducer positions, that a smaller focus induced reduced activation and that lateral sonications produced an ipsilateral bias in the activated areas. Additionally, a previous study from our group that combined functional ultrasound with displacement imaging has demonstrated that the displaced area of brain tissue is larger than the area of the FUS focal spot in the brain and that FUS neuromodulation along the midline can cause bilateral hemodynamic responses [44]. However, it is also conceivable that the lack of focality was caused by off-target effects, as previous work from our group has indicated that intracranial reverberations could generate high pressure amplitudes outside of the intended focal target [11]. Further studies that combine fUSI with electrical or optical recordings of neuronal activity in multiple brain regions are needed to fully elucidate the short-term activation patterns generated by FUS neuromodulation in the brain.

Subcortical activation was inconsistent during transcranial fUSI acquisitions while reliable activation was observed in the cortex. In mice implanted with an ultrasound-transparent cranial window, we found more robust activation in subcortical regions and less pronounced cortical activation. These results could be explained by the effects of temperature on neuronal activation since the brain temperature was decreased in mice with a cranial window, especially in cortical areas. Thermocouple measurements revealed a temperature of 31.6 °C in the cortex and 33.6 °C in the thalamus in the cranial window condition (Supplementary Figure 4), whereas brain temperature under similar anesthetic conditions without a cranial window has been reported by others to be 34.6 °C or 35.4 °C, respectively [49]. In addition, FUS can cause skull heating due to the acoustic properties of the skull, which results in a local temperature increase below the skull surface [50,51]. Thermocouple measurements directly under the skull showed temperature increases of up to 3.58 °C with the highest pressure used in the transcranial conditions (Supplementary Figure 3 b). Although we do not expect these temperature increases to cause damage due to the cooling effect of the ultrasound gel on the mouse skull, it has been shown repeatedly that temperature can influence neuronal activation and cerebral blood flow [52–55]. Specifically, temperature decreases induce lower cerebral blood flow, while the effects on neuronal activity are mostly described as excitatory [52,54]. Both mechanisms as well as skull heating likely influence the results presented here and could explain the differences between the activation profiles in the transcranial and cranial window conditions. Furthermore, lower temperature decreases the activity of mechanosensitive channels like Piezo1 [56] and TRPP2 [57] which have been shown to excite neurons during FUS neuromodulation [58]. Further investigation of the effects of temperature in the context of fUSI and during FUS neuromodulation is therefore warranted to advance our understanding and refine the application of both techniques. In addition, the stronger cortical activation and less pronounced subcortical response in the transcranial condition compared to animals with a cranial window should caution authors to directly translate results between both cases and carefully adjust relevant parameters like the cranial temperature.

Interestingly, we were able to identify negatively correlated regions with stimulation-associated decreases in CBV in mice implanted with a cranial window, which could be connected to decreases in neuronal activity [59]. However, a variety of different mechanisms like a ‘steal-phenomenon’ [60] in the vicinity of active regions or vasoconstriction independent of neuronal activity [61] have been proposed. The absence of negatively correlated regions in the transcranial condition could indicate a lower sensitivity in transcranial imaging. The association of higher frequency with more negatively correlated pixels indicates an effect of frequency or focal size on the hemodynamic responses during FUS that warrants further investigation.

Limitations of this study include the inability to rule out thermal effects on neuronal signals and vascular responses in the brain, especially given that the highest pressures induced temperature increases > 2 °C in the brain and under the skull. Despite this, fUSI was able to pick up FUS-induced hemodynamic changes even in lower pressures that caused only minor temperature increases (< 1 °C). Another limitation comes from the variation in the imaging planes used to study the effects of FUS with fUSI, especially in the transcranial condition. This does not allow the precise identification of possible pathways activated by FUS neuromodulation. However, the fact that we were able to nevertheless find the demonstrated relationships with FUS pressure and lateral bias of responses could suggest robust effects on wider brain networks. Another limitation is that auditory artifacts could not be completely ruled out as a potential confounder in the hemodynamic activations. This concern is particularly relevant in light of recent research indicating that auditory confounds are a primary factor driving FUS neuromodulation effects in humans [62]. However, the absence of activation in an auditory sham condition, along with the dose-dependent effects and lateralized responses observed with FUS, strongly indicates that the mechanism of action is not predominantly auditory. To conclusively eliminate auditory artifacts, future studies will entail a transgenic, conditionally deafened mouse model, as described by Guo et al. [63].

## Conclusion

This study introduced a system for FUS neuromodulation that allows simultaneous online monitoring of hemodynamic responses with fUSI *in vivo*. We show that fUSI can capture region-dependent responses to FUS neuromodulation and displays stronger responses in higher-pressure conditions. Our approach allowed for transcranial imaging of FUS neuromodulation-induced changes with a large field of view in rodents, which could help in studying the immediate to mid-term effects of FUS neuromodulation, especially in the context of network activation patterns.

## Data Availability

Datasets and executable codes can be shared by the corresponding author upon reasonable request.

## CRediT authorship contribution statement

**Jonas Bendig:** Conceptualization, Formal analysis, Investigation, Methodology, Project administration, Visualization, Writing – original draft, Writing – review & editing. **Christian Aurup:** Conceptualization, Formal analysis, Investigation, Methodology, Project administration, Software, Visualization, Writing – original draft, Writing – review & editing. **Samuel G. Blackman:** Software, Visualization, Writing – review & editing. **Erica P. McCune:** Visualization, Writing – review & editing. **Seongyeon Kim:** Project administration, Software, Writing – review & editing. **Elisa E. Konofagou:** Conceptualization, Funding acquisition, Methodology, Supervision, Writing – original draft, Writing – review & editing.

## Supporting information

Supplementary Video 1. CBV with FUS artifact in a craniotomized mouse

## Acknowledgments

This work was supported by the National Institutes of Health (Award numbers: 5R01EB02757 6-04 and 5R01AG03896 1-10) and Jonas Bendig was partially supported by the Thiemann Foundation and the German Research Foundation. Additionally, we would like to thank Dr. Stephen Lee for his crucial advice and support during the whole project.

