## Supplementary Figures for "Transcranial Functional Ultrasound Imaging Detects Focused Ultrasound Neuromodulation Induced Hemodynamic Changes *In Vivo*"

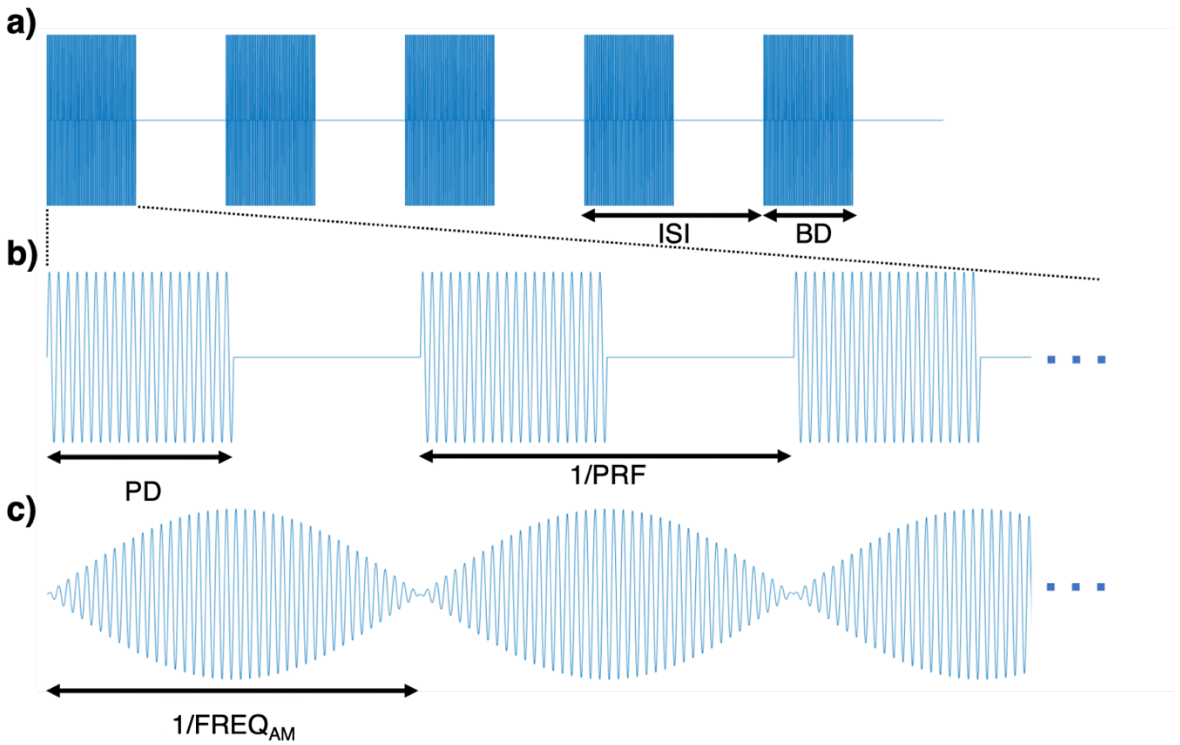


**Supplementary Figure 1.** Pulsing scheme and comparison of square envelope with amplitude modulation. The FUS stimulus consisted of several bursts of ultrasound **(a)** with a burst duration (BD) of 150 ms. The inter-stimulus interval (ISI) delineates the time between the onset of one burst and a consecutive burst (ISI = 1/FREQ). For the classical square wave envelope **(b)**, a burst consists of interrupted ultrasound pulses with a pulse duration (PD) that is determined by the pulse repetition frequency (PRF) and the duty cycle (DC). The amplitude of the underlying ultrasound wave does not change over the PD. In the scheme used in this study **(c)**, the amplitude of the underlying ultrasound wave is modulated with a sinusoidal wave with a frequency (FREQ) of 1 kHz.

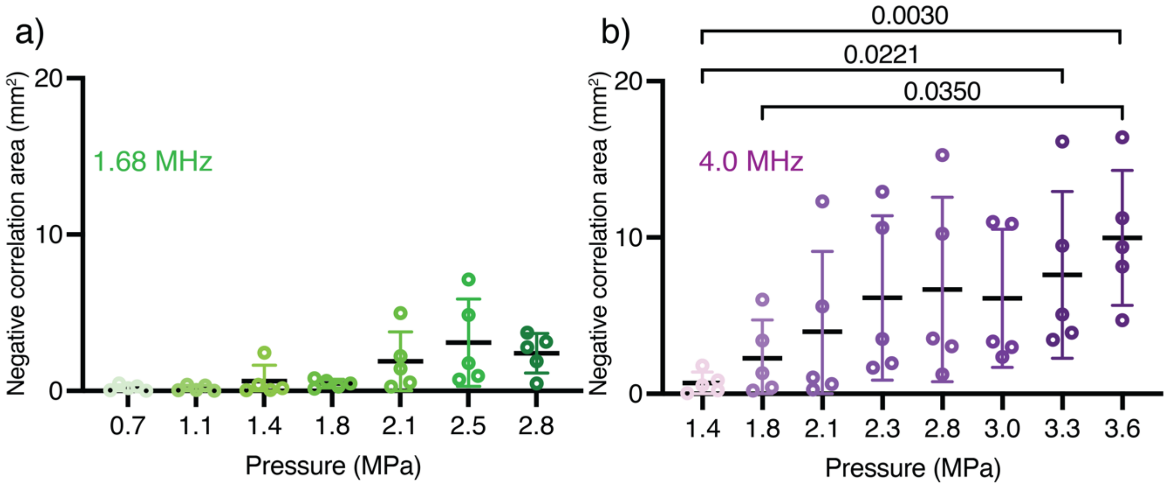


**Supplementary Figure 2.** Areas with significantly negative correlation values in the cranial window condition (corresponding to Figure 4). Graphs show the area of all significant negatively correlated pixels (Spearman's $\rho < -0.19$) in the 1.68 MHz condition (**a**, n = 5 paired trials in 5 mice, median AP direction of imaging planes: -1.5 mm from Bregma, Range = -1.4 mm to -1.9 mm; Friedman test: $p = 0.0104$) and the 4.0 MHz condition (**b**, n = 5 paired trials in 5 mice, median AP direction of imaging planes: -1.5 mm from Bregma, Range = -1.4 mm to -2.0 mm; Friedman test: $p = 0.0003$). Data is depicted as individual observations (circles) with mean (central line) and standard deviations (whiskers). Dunn's test (corrected for multiple comparisons) was used for post-hoc comparisons after significant Friedman tests. For the 1.68 Mhz condition, post-hoc tests showed no significance for individual comparisons.

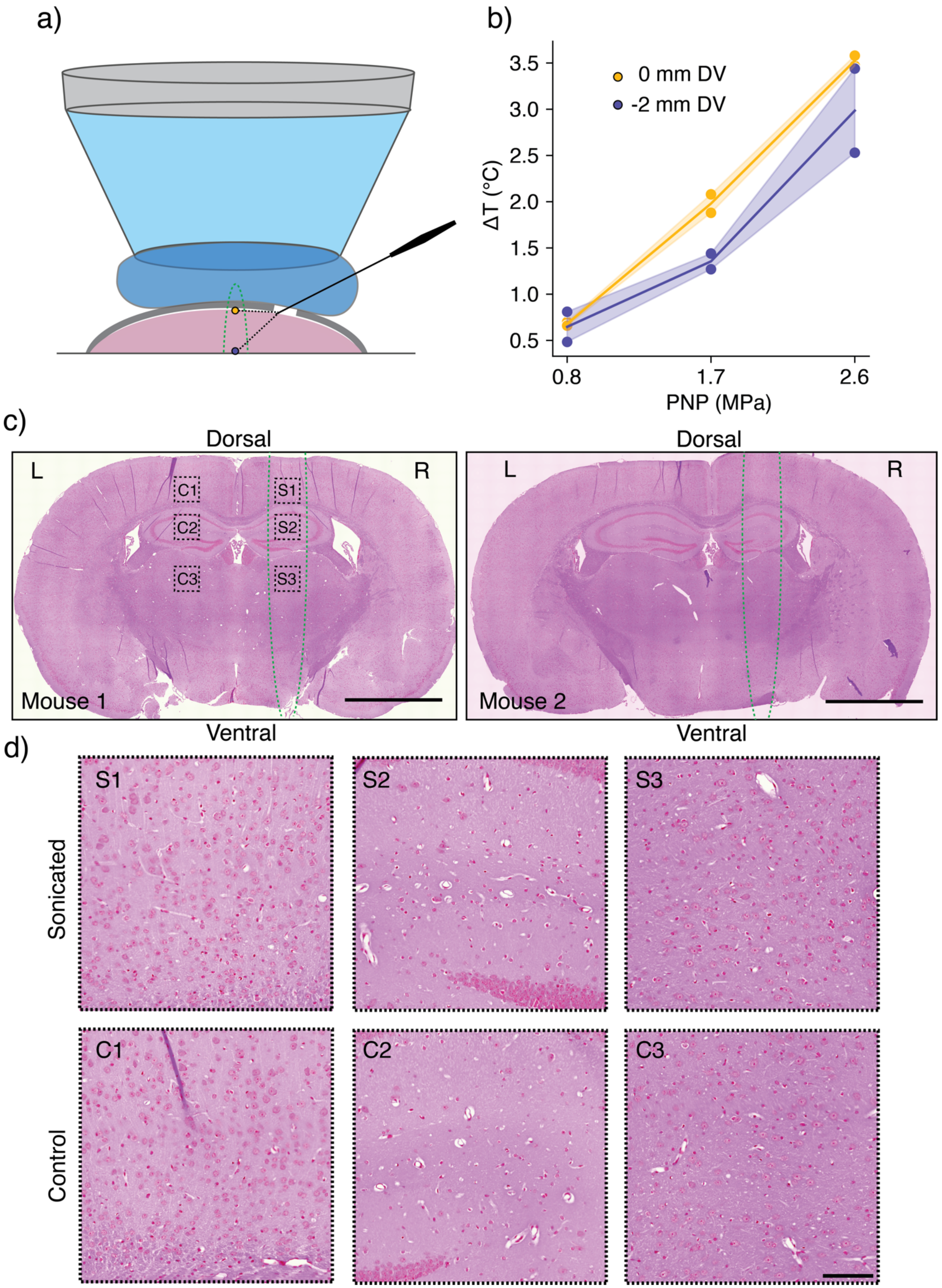
a)
b)
0 mm DV
-2 mm DV
ΔT (°C)
3.5
3.0
2.5
2.0
1.5
1.0
0.5
0.8
1.7
2.6
PNP (MPa)
c)
Dorsal
L
R
C1
C2
C3
S1
S2
S3
Mouse 1
Ventral
Dorsal
L
R
Mouse 2
Ventral
d)
Sonicated
S1
S2
S3
Control
C1
C2
C3

21

**Supplementary Figure 3.** Safety and temperature measurements. A schematic of the setup for temperature measurements in the brain is shown in (**a**). A thermocouple (HYP1, T-type needle thermocouple, 0.25 mm diameter, Omega Engineering, Norwalk, CT) was inserted into the brain of anesthetized mice (n = 2) with a micromanipulator through a small burr hole in the skull. Temperature measurements during FUS were performed at two locations (DV -2 mm from the brain surface and at DV 0 mm, directly under the skull. Temperature measurements corrected for viscous heating artifact show strong temperature increases in the highest pressure condition **(b)**. Additionally, safety was assessed in n = 2 separate animals 24 h hours after transcranial sonication with the highest FUS pressure (PNP = 2.6 MPa). H&E staining showed no signs of tissue damage or red blood cell extravasation **(c/d).** The estimated focal position is overlaid with a green dotted line in **c.** Panels in **d** show magnifications of the correspon**ding** regions (cortex, hippocampus, thalamus) from **c** on the sonicated **(S)** and unsonicated **(C)** hemispheres for comparison. Sonications were performed at ML -1 mm, AP -1.5 mm from Bregma. Scale bars 2 mm in **c**; 50 µm in **d**. Abbreviations: AP – anteroposterior; C – control; DV – dorsoventral, L – left; R – right; S – sonicated. Points represent individual measurements in, solid line represents the mean and the shaded area the 95 % confidence interval (b).

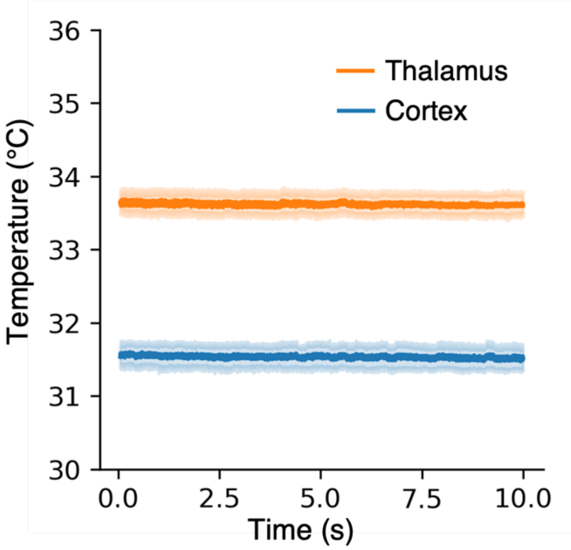


**Supplementary Figure 4.** Temperature in different areas under the cranial window. 10 short thermocouple measurements (10 s each) were performed evenly spaced over a period of 30 min in the cortex and the thalamus ($n_{animals}$ = 1). During the whole period, the acoustic coupling cone and transducer were coupled to the mouse head with ultrasound gel as described in the methods. The temperature measurements were performed without FUS neuromodulation or fUSI. Measurements were stable over time during all measurements for both locations. Shaded areas depict 95 % confidence intervals. Mean temperatures: Cortex = 31.53 °C (±0.24 °C), Thalamus = 33.61 °C (±0.26 °C).
